# The role of temporal coherence and temporal predictability in the build-up of auditory grouping

**DOI:** 10.1101/2021.06.27.449898

**Authors:** Joseph Sollini, Katarina C Poole, Dominic Blauth-Muszkowski, Jennifer K. Bizley

**Affiliations:** Hearing Sciences, Mental Health and Clinical Neurosciences, School of Medicine, University of Nottingham, England, United Kingdom; The Ear Institute, University College London, London, England, United Kingdom

## Abstract

The cochlea decomposes sounds into separate frequency channels, from which the auditory brain must reconstruct the auditory scene. To do this the auditory system must make decisions about which frequency information should be grouped together, and which should remain distinct. Two key cues for grouping are temporal coherence, resulting from coherent changes in power across frequency, and temporal predictability, resulting from regular or predictable changes over time. To test how these cues contribute to the construction of a sound scene we present listeners with a range of precursor sounds, which act to prime the auditory system by providing information about each sounds structure, followed by a fixed masker in which participants were required to detect the presence of an embedded tone. By manipulating temporal coherence and/or temporal predictability in the precursor we assess how prior sound exposure influences subsequent auditory grouping. In Experiment 1, we measure the contribution of temporal predictability by presenting temporally regular or jittered precursors, and temporal coherence by using either narrow or broadband sounds, demonstrating that both independently contribute to masking/unmasking. In Experiment 2, we measure the relative impact of temporal coherence and temporal predictability and ask whether the influence of each in the precursor signifies an enhancement or disruption of unmasking. We observed that interfering precursors produced the largest changes to thresholds.

## Introduction

It is often hard to fathom how the auditory system is able to turn a simple two-dimensional signal, consisting of variations in pressure over time, into the complex sound scene we perceive. The auditory system uses cues to group together information arising from separate sources, or ‘objects’^1-4^ and segregate information arising from distinct sources^5^. Here we focus on two important cues and their role in establishing auditory grouping, temporal coherence and temporal predictability. Sound elements from a common source will often fluctuate in power simultaneously. These synchronous changes in amplitude across-frequency serve as a critical grouping cue, useful for sound detection^6,7^ and speech processing^8^, and can be referred to as either comodulation^9^ or temporal coherence^4^ (Note: as the amplitude fluctuates the availability of other cues will become available/unavailable and temporal coherence refers to the fluctuation in these cues over time also). A sound’s statistics can be stationary over time (for example a pure tone of fixed frequency) or vary over a defined range, such as natural sound textures^10^. Reliability in these statistics over time also improves the segregation of auditory objects^11-13^. An extreme example of reliable temporal statistics (referred to here as “temporal predictability”) are sound patterns where the temporal structure is repetitive (or regular) over time and, hence, it’s statistics vary over a very narrow range^14^.

The auditory system is excellent at gathering information over time to improve perception, this process of accumulating knowledge about a sound is often referred to as the “build-up” period. Build-up has been observed in a range of seemingly disparate auditory behaviours, such as listening to speech-in-noise^15^, speech in speech^11^, auditory stream formation^16^, the discrimination of sound textures^10^, comodulation masking release^17^, the precedence effect^12^ and transitions to new spectrotemporal sound statistics^18^. Generally speaking, the longer the build-up period, the greater the behavioural benefit^10,12,19-21^, although, this benefit will asymptote and can subsequently decline due to inattention^22^. The optimal build-up period can be thought of as the shortest duration of time over which performance begins to asymptote, i.e. when additional build-up offers very little behavioural advantage. The optimal build-up period varies depending on the task, but appears to be between 0.4-2.5 seconds^10,12,15,19^, though further modest gains can be made for more than a minute^16,23^. In the case of “predictable” sounds, i.e. repeating spectrotemporal structure, this variation can be explained by the duration or period of the sound statistics themselves, i.e. the longer a repeating sequence the more time is required to learn that sequence^18^.

Once an object has been built-up it can take several seconds for its influence to decay^16,24^ unless object formation is reset by abrupt changes in an ongoing sound, e.g. a frequency, level or spatial deviation in one of the objects^23,25-28^. Overall, the existing literature suggests that build-up is a high-level evidence accumulation process^21^ that influences auditory processing. It can do this by either adapting to or enhancing the important features within an object and operates on a range of grouping cues. Establishing the structure in an auditory scene then serves a range of core auditory abilities, such as segregation^11,16^, detection^13,29^ or discrimination^10^. It is not clear whether build-up is one general process or whether independent build-up mechanisms subserve different grouping cues. To better understand this, it is necessary to investigate interactions between different grouping cues in order to measure their influence upon one another.

Here, we investigate how temporal predictability and temporal coherence interact with one another by manipulating the cues available during the build-up period. Given the reasonably long-lasting effects of build-up (up to 1 second) these interactions should have important consequences on the perception of subsequent sounds. Indeed, grouping processes have been shown to effect auditory stream segregation^16,30^ and also sound detection^31-33^. We chose to employ a simple tone in background sound detection task in which the likelihood of detecting the tone was influenced by the ability of listeners to successfully segregate the tone and masker. This provides an indirect but objective measure of how the listener is grouping the sound mixture. Here, increases in tone detection threshold indicate an impaired ability to represent both sounds (masker and tone) separately, while decreases in detection threshold suggest that the masker and tone are perceptually better segregated. We sought to promote perceptual segregation with the use of either temporal coherence and/or temporal predictability as a grouping cue during the build-up period. By varying the background sound statistics during build-up, while providing both cues later in the background (when the tone was presented) we were able to assess how these statistics influenced subsequent grouping. Using this approach, we were able to show that manipulating temporal predictability and coherence in the build-up can significantly affect the auditory systems ability to make use of these grouping cues. Our results demonstrate the importance of the build-up period on organising an auditory scene and thus the subsequent perception of auditory objects.

## Materials and methods

### Participants

All participants were aged between 19 and 25 and none reported long-term or current problems with their hearing. Experimental procedures were carried out in accordance with the protocols approved by the research ethics committee of University College London (reference 3866/002) and written informed consent was provided by each participant. Participants were given a short training session (usually lasting 15-20 minutes) to familiarise themselves with the experimental setup and the task itself before starting.

## Stimulus and apparatus

### Apparatus

Participants were seated in a double-walled sound attenuating booth (IAC, Winchester, UK), wearing headphones (Sennheiser HD 600), in front of a computer monitor and mouse. Sound was delivered through the headphones via a preamplifier connected to a signal generator (Tucker Davis Technology, RP2) which was in turn controlled by custom written software (MATLAB and TDT’s RPvdsEX). Participants registered responses to sounds via a custom designed graphical user interface (MATLAB) through mouse clicks at the end of each trial.

Sound stimuli were generated *de novo* for each trial using MATLAB and loaded onto the signal generator at the start of each trial. All sounds were generated and presented with a ∼48kHz sampling rate.

### Signal

Participants were required to detect a short (25ms duration, 12ms cos^2^ ramp) pure tone signal (1 kHz) that was embedded in a masker. Tone signals were presented in the trough between peaks 13 and 14 of the background (see below), meaning they were centred, temporally, in the middle of the masker (e.g. Fig. 1A). To measure tone detection thresholds, tones were varied in sound level according to an adaptive staircase rule (see procedure).

**Fig. 1.**
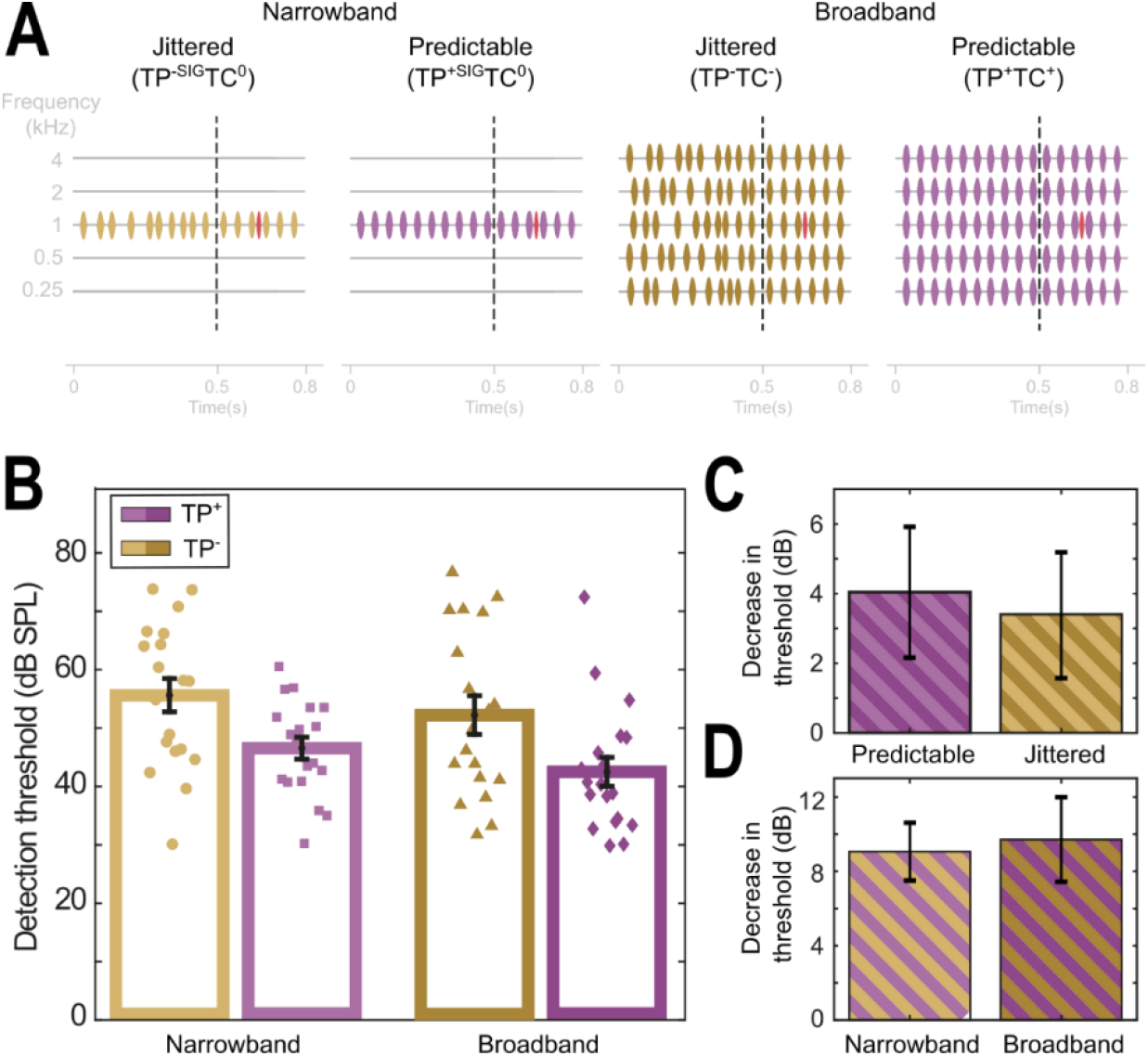
Temporal coherence and temporal predictability during build-up significantly effect subsequent detectability, when the two cues are not in conflict there is no evidence of an interaction. A) Stimulus schematic. Each stimulus was composed of a background (purple/green) and a pure tone signal (red). The backgrounds were a series of pure tone pips presented in either: i) Left two panels) a narrowband (TC^0^ notation) or ii) Right two panels) broadband (TP^+/-^ notation, 5 × 1 octave spaced tones centred at 1kHz) configuration. Each background was composed of a pre-cursor (left of the vertical dashed line) and a masker (right of the vertical dashed line). The pre-cursor could either promote temporal predictability (purple and TP+ notations, including TP^+SIG^) or disrupt temporal predictability (green and TP^-^ notation, including TP^-SIG^). B) Pure-tone detection thresholds for each condition in (A). C) The reduction in masking attributable to bandwidening was similar across the two temporal predictability conditions (Predictable = TP^+SIG^TC^0^ vs TP^+SIG^TC^0^ and Random = TP^-SIG^TC^0^ vs TP^-^TC^-^, Predictable = ∼4 dB, Random = ∼3.4 dB). D) The reduction in masking attributable to adding temporal predictability (in the precursor) was also similar (Narrowband = TP^+SIG^TC^0^ vs TP^-SIG^TC^0^ and Broadband = TP^+^TC^+^ vs TP^-^TC^-^, Narrowband: ∼9.1 dB, Broadband: ∼9.7 dB).

### Background stimuli

To create the backgrounds, modulated pure tones (800ms duration) were created with a random phase at the necessary frequencies for the narrowband (1kHz, 65dB SPL) and broadband conditions (1kHz as before but also at: 0.25, 0.5, 2 and 4kHz, termed ‘flanker bands’ where each was calibrated to be 65dB SPL). The background was composed of two periods of time; the precursor (the first 500ms) and the masker (the next 300ms). In both experiments the masker was always temporally predictable (individual “pip” envelopes generated using ½ a cycle of a 40 Hz cosine which were then positioned at regular intervals at a 20Hz rate). Temporal predictability, (TP), and temporal coherence (TC) in the precursor were varied in the two experiments. In Experiment 1 the precursor and masker could be broadband (TP^+^TC^+^ and TP^-^TC^-^, see Fig. 1) or narrowband (TP^+^TC^0^ and TP^-^TC^0^) where the precursor could be either predictable (TP^+^TC^+^ and TP^+^TC^0^) or random (TP^-^TC^+^ and TP^-^TC^0^, each pip onset within the precursor was adjusted by a uniformly sampled random amount of time between the bounds ±25ms).

In Experiment 2 the precursor and masker were always broadband and the temporal predictability and coherence of the signal band (1kHz frequency band) and off-frequency components (0.25, 0.5, 2 and 4kHz) were manipulated yielding 4 conditions with different precursors (see Fig. 2): temporally regular across all frequency bands (TP^+^TC^+^), jittered within the signal band (TP^-SIG^TC^-^), the flanker bands (TP^+SIG^TC^-^), or both (TP^-^TC^-^), and an additional ‘no-build up’, or neutral TP^0^, condition which just contained the masker with no precursor. Schematics of the stimuli in each experiment are included in Figure 1A and 2A.

**Fig. 2.**
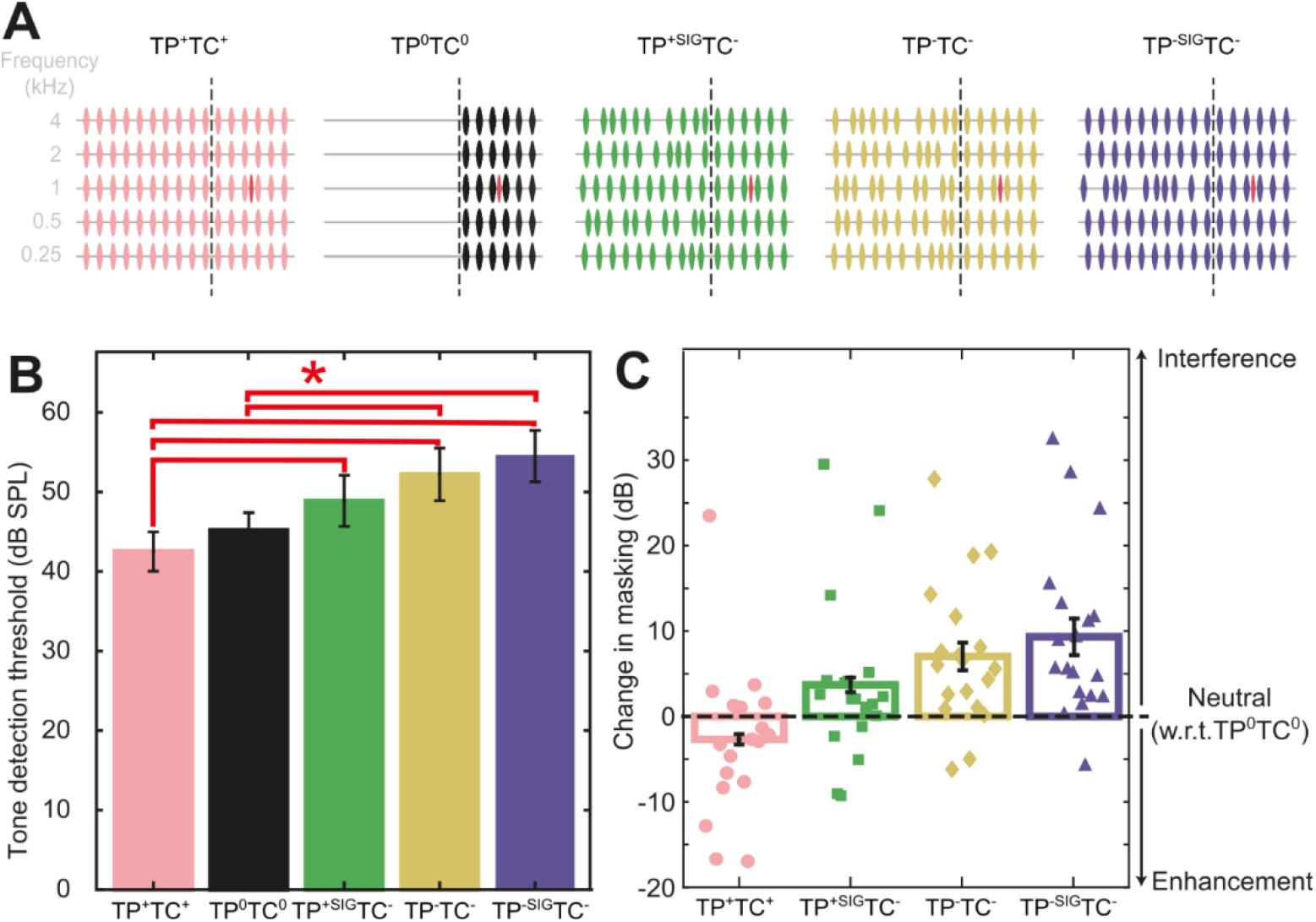
Discontinuity in sound statistics (temporal coherence and/or predictability) causes interference in the grouping process. A) Stimulus schematic. Five conditions were contrasted: a neutral condition (i.e. no build-up: TP^0^TC^0^) and four where the build-up was varied (i.e. TP^+^TC^+^ = both temporal coherence and predictability, TP^+SIG^TC^-^ = Temporal predictability in the signal band only and random temporal coherence, TP^-^TP^-^ = Unpredictable (random) and no temporal coherence and TP^-SIG^TP^-^ = Temporal predictability and coherence in the flanking bands but random structure in the signal band). B) Tone detection thresholds for each condition. C) The aim of these manipulations was to measure the relative contribution of “enhancement” (decrease in threshold w.r.t. the neutral condition) and “interference” (relative increase in threshold w.r.t. the neutral condition). Temporal predictability and coherence in the build-up tended toward enhancing the tone (though did not reach significance), whereas, random structure in the signal band (TP^-SIG^TC^-^) or in all frequency bands (TP^-^TC^-^) produces significant interference.

## Testing protocol

### Staircase procedure

An adaptive staircase procedure was used to vary signal level between trials. A 3up 1down design was used with three sequential rules for varying the change in sound level: rule 1 = 3 reversals with a 6 dB step size, rule 2 = 3 reversals with a 2 dB step size and rule 3 = 4 reversals with a 1dB step size. The tone signal level started at 65 dB SPL for each block.

### General procedure

Tone detection threshold for each of the seven backgrounds were measured in blocks and 3 repeats of each threshold were measured. The intention was for that for each repeat, the order of the blocks would be re-randomised. Owing to an experimenter error this was only performed for 8 of the 19 participants. Nonetheless, statistical comparison of the mean thresholds across the three repeats for each condition for the two groups (i.e. randomised vs. same order participants) demonstrated there was no significant effect of the order within a repeat cycle between the two groups (paired t-test, uncorrected for multiple comparisons to add confidence, p>0.05 for all conditions). Within each block a 3-interval forced choice design was used to probe tone signal detection. On a given trial 3 identical backgrounds, presented consecutively, were presented with a 1 second interval between each. Within one of these intervals a tone signal was added, and the participant was asked to identify the interval containing the tone using a mouse click on a response screen.

### Data analysis

Data analysis was performed in MATLAB. For each participant and testing block the threshold was calculated as the mean signal sound level across the final 4 reversals and averaged across the three repeats of each condition. Statistical analysis was performed in SPSS (IBM SPSS Statistics 27).

## Results

### Temporal predictability and temporal coherence can both independently influence grouping

To test the role of temporal predictability and temporal coherence in build-up we employed a simple tone in background stimulus design. In brief, participants (n=19) were presented the same background 3 times in consecutive intervals (0.8 second duration, 1 second inter-stimulus interval), in one of these intervals a tone was also presented, and the participant had to identify this interval (i.e. 3-interval forced choice design). From trial-to-trial the sound level of a target tone was varied with a 3-down, 1-up adaptive staircase in blocks to yield a threshold for each block (the mean tone level across the last 4 reversals) and then for each condition (the mean of all 3 blocks/repeats for that condition).

In Experiment 1, we compared the impact of manipulating temporal coherence and predictability in the precursor (Fig 1A). To do this, four backgrounds were compared in a 2×2 design. The effect of across-channel temporal coherence was tested by altering the bandwidth, comparing a narrowband (1kHz tone pips) and a broadband configuration (5 frequency complex between 0.25 and 4kHz with 1 octave spacing). In addition, the effect of temporal predictability was tested by varying the stimulus statistics in the build-up (i.e. precursor) while maintaining identical stimulus statistics in the masker portion of the background, across comparisons (500-800ms). The statistics in the masker period were always regular and were narrowband or broadband to match the build-up, but the temporal structure in each frequency channel during the build-up could either be regular (i.e. temporal predictability, Fig. 1A) or jittered (unpredictable, Fig1A). Note that for the broadband condition the temporally predictable configurations were also temporally coherent across frequency. In total Experiment 1 had four conditions: two narrowband conditions that allowed us to measure the effect of temporal predictability in the absence of across-channel temporal coherence, one condition was predictable during the build-up (TP^+SIG^TC^0^, Fig. 1A) and the other jittered (TP^-SIG^TC^0^). The two broadband conditions allowed the assessment of varying temporal predictability and coherence during build-up: one with temporal predictability and a temporally coherent build-up (TP^+^TC^+^) and one without (TP^-^TC^-^, though note that in both cases the masker did have temporal coherence and predictability).

Central to our hypothesis was the assumption that a lower (better) tone detection threshold would result when listeners were better able to segregate the masker from the competing tone.

Comodulation Masking Release is a well-known auditory phenomena where adding temporally coherent off-frequency maskers paradoxically results in a lower thresholds^7^, thought to result from perceptual grouping of the signal and flanking channels^32,34-36^. We predicted that thresholds should be lowest in the TP^+^TC^+^ case, where the broadband masker would enable masking release and the build-up period contained a temporally predictable and temporally coherent structure matching that in the masker. We predicted that if temporal predictability plays a role in object formation both temporally predictable conditions should yield lower thresholds than the jittered conditions (i.e.TP^+^TC^+^ < TP^-^TC^-^ and TP^+SIG^TC^0^ < TP^-SIG^TC^0^).

Varying either bandwidth or temporal predictability had a significant effect on subsequent tone detection thresholds (Fig 1B). Adding across-channel temporal coherence in the masker produced significant unmasking of the tone target, irrespective of temporal structure in the build-up (µ = 3.72 dB, SD = 7.67dB). Altering the temporal structure, by transitioning from a jittered to a temporally regular modulation rate, whether broadband or narrowband (Fig. 2A), produced a large reduction (µ = 9.39 dB, SD = 7.13dB) in tone detection thresholds. These findings were confirmed with a two-way repeated measures ANOVA with main effects of adding bandwidth and predictability (predictability: F(1,18) = 28.45, p = 0.000045, bandwidth: F(1,18) = 5.171, p = 0.035). Improvements in threshold produced by unmasking due to bandwidth (Fig. 1C) and predictability (Fig. 1D) remained consistent across the other manipulation, confirmed by the non-significant significant interaction term (predictability x bandwidth: F(1,18) = 0.151, p = 0.702). The benefit observed by adding additional frequency bands was identical in magnitude for both jittered and predictable maskers, suggesting that varying temporal predictability during build-up did not disrupt the across-channel benefit of temporal coherence in the masker (Fig. 1C). In addition, the benefit of having temporal predictability within the precursor was also similar regardless of the bandwidth (Fig. 1D). Overall, predictability produced significantly larger reductions in threshold than band-widening (µ = 9.39 vs 3.72dB, respectively, paired t-test, t(36)= 2.3598, p = 0.0238).

### Build-up critically influences the subsequent use of both temporal predictability and coherence cues

In Experiment 1 it was difficult to tease apart the relative contribution of temporal predictability and temporal coherence to subsequent perception. For example, providing predictability and coherence in the build-up (TP^+^TC^+^) could facilitate grouping of the on-frequency channel into the background and, hence, enhance tone detection. Alternatively, disrupting predictability and coherence during the build-up (TP^-^TC^-^) could interfere with subsequent grouping and tone detection. When comparing these conditions, and observing a difference, it is unclear whether we are measuring either one or both. To allow quantification of the relative enhancement/interference on subsequent performance we introduced a “neutral” condition where the build-up period had been removed (Fig 2A, TP^0^TC^0^). This allowed us to determine the relative direction of change in performance that was elicited by the inclusion of each respective precursor.

Experiment 1 suggests that temporal predictability and coherence are independent processes, as the size of one was not significantly varied when changing the parameters of the other (and confirmed by the non-significant interaction term in the ANOVA). However, in this experiment the jittered precursor transitioned to a regular sequence of tones in the masker. These conditions favour treating the precursor and masker as independent from one another, as this transition is likely to be interpreted as the start of a new auditory object. In contrast in the predictable case the continuity in statistics will cause the precursor and masker to be grouped together. For the broadband case the cues are consistent across all frequencies. We next sought to test how temporal coherence and predictability cues interact when put into conflict (from precursor to masker), by independently modulating the temporal structure of the flanking and signal bands in the build-up period.

In Experiment 2 we generated two additional conditions, each designed to promote different grouping configurations, while disrupting the grouping of flanking and signal bands during the precursor. The first promoted the use of temporal predictability to group just the signal band, by providing continuous regularity in the signal channel from precursor to masker, while jittering the timing of the flanking bands in the precursor (TP^+SIG^TC^-^, Fig. 2A). This manipulation aimed to block the use of temporal coherence in the masker but allow the use of temporal predictability in the signal band. The second sought to block both the use of temporal coherence (across-channels) and temporal predictability (within the signal band, TP^-SIG^TC^-^).

A significant effect of varying the masker properties during build-up was found across the 5 conditions (RM-ANOVA of all five conditions, n = 19, F(4,18) = 11.729, p = 0.0000002). Tone detection thresholds were lowest when the build-up contained identical sound statistics to the masker period (TP^+^TC^+^, µ = 42.5dB SPL) suggesting an enhancement relative to the neutral condition (TP^0^TC^0^, µ = 45.12dB SPL, 2.62dB difference), however, post-hoc test revealed this was not significant (paired t-test, Sidak correction, p = 0.897). By contrast, conditions with jitter in the build-up produced interference, relative to the neutral condition (Fig. 2B: TP^-^TC^-^, µ = 52.21, TP^-SIG^TC^--^, µ = 54.5, and TP^+SIG^TC^-^ µ = 48.89dB SPL, or 7.1, 9.38 and 3.77dB increases respectively). Post-hoc tests revealed interference was significant when all frequency channels were jittered (TP^-^TC^-^) or the signal band was jittered (TP^-SIG^TC^-^) but not when the flanking bands were jittered and the signal band had temporal predictability (TP^+SIG^TC^-^, paired t-test, Sidak correction, p = 0.019, 0.007, 0. 699 for TP^-^TC^-^, TP^-SIG^TC^-^ and TP^+SIG^TC^-^ respectively). While neither the TP^+^TC^+^ or TP^+SIG^TC^-^ conditions were significantly different from the neutral condition (TP^0^TC^0^), they were significantly different from one another (6.39dB difference, paired t-test, Sidak correction, p = 0.049), suggesting they produce small changes from the neutral condition that only become significant when contrasted. In addition, TP^+^TC^+^ thresholds were significantly lower than the other two unpredictable conditions (paired t-test, Sidak correction, p = 0.00031 and 0.0046, TP^+^TC^+^ vs TP^-^TC^-^ and TP^-SIG^TC^-^ respectively). Together these results demonstrate that the preceding sound statistics can significantly affect subsequent sound in noise detection. Interference is largest when the signal band’s temporal statistics are unpredictable (i.e. TP^-^TC^-^ and TP^-SIG^TC conditions). Smaller changes are observed when temporal predictability in the build-up is present (TP^+SIG^TC^-^, i.e. when only disrupting temporal coherence in the flanking bands) or providing both cues (TP^+^TC^+^).

### Build-up can be used to block the subsequent use of temporal coherence cues

In Experiment 2, conditions that discouraged grouping of the signal and flanking bands during build-up, i.e. TP^+SIG^TC^-^ and TP^-SIG^TC^-^, appeared to produce comparable thresholds to those observed in the narrowband conditions in Experiment 1 (Fig. 1B). As the same subjects completed Experiment 1 and 2 we were able to directly compare thresholds across both. Therefore, thresholds for the narrowband conditions in Experiment 1 (TP^+^TC^0^ and TP^-^TC^0^) were directly compared to the relevant conditions in Experiment 2 (TP^+SIG^TC^-^ and TP^+SIG^TC^-^, respectively). We performed a two-way repeated measures ANOVA with within-subject factors of bandwidth and predictability. This yielded a significant effect of predictability (F(1,19) = 35.7, p = 0.000012, p<0.05 after Bonferonni correction) but not of bandwidth (F(1,19) = 7.286, p = 0.722). In addition, no interaction was found (predictability*bandwidth, F(1,19) = 2.33, p = 0.144). These results suggest that the benefits of bandwidening observed in Experiment 1 (Fig. 1C) could be blocked by the addition of incoherent flanking/signal band envelopes in the build-up period.

## Discussion

We employed a simple paradigm to assess how manipulating grouping cues during ‘build-up’ influenced subsequent sound segregation, as assessed by a tone detection task. We found that those grouping cues available during build-up critically shaped subsequent perception. Eliminating predictability during build-up significantly hampered sound segregation, and elevated tone detection thresholds, relative to conditions with temporal predictability in the signal band. In addition, disrupting temporal coherence between signal and flanker bands during the build-up also produced significant increases in thresholds. Thus, these results demonstrate the influence of early sound statistics on subsequent sound perception.

The general pattern of results observed here were consistent with the idea that object formation during build-up is extremely influential to subsequent perception, at least within the relatively recent context. Conditions that discouraged grouping of the signal and flanking bands, i.e. TP^+SIG^TC^-^ and TP^-SIG^TC^-^, produced thresholds comparable to those observed in the narrowband conditions. For example, build-up could be used to block the subsequent use of temporal coherence cues (last paragraph of the results). Previous work demonstrating that surrounding a period of temporal coherence with random temporal structure (similar to TP^-^TC^-^) disrupts the benefits of temporal coherence^37,38^. One could argue that a lack of temporal coherence during build-up (TC^-^) indicated to the auditory system that the signal and flanking bands should not be grouped (as there is no evidence they should be grouped). However, this is insufficient to explain the results in Experiment 1 where the temporal coherence benefit was comparable for the jittered and predictable comparisons (see Fig. 1C). Here, the lack of interaction demonstrates that bandwidening produced the same effect size regardless of the coupling (TC^+^)/decoupling (TC^-^) of channels during build-up. This suggests that the ability to utilize across-channel grouping cues in the masker was not blocked by their absence in the build-up period. In Experiment 2, thresholds were higher for the TP^-SIG^TC^-^ than for the TP^-^TC^-^, suggesting that if an object had been established during the precursor that did not include both signal and flanker bands (through temporal predictability in flanker or signal but not both) then this could block the subsequent use of temporal coherence in the masker. This suggests that object permanence of temporal stability is not overridden by temporal coherence, at least for the limited set of parameters used here. This might be an exception to the temporal coherence model, i.e. “that stream formation depends primarily on temporal coherence between responses that encode various features of a sound source”^4^, as object permanence can block the use of temporal coherence to form streams.

It should be noted that the masker period had a very short build-up (between the offset of the pre-cursor and the onset of the tone) of 187.5 milliseconds. It may be that the large change in sound statistics is sufficient to “reset” build-up, similar to resetting of stream segregation^25,26^, and there is time for temporal coherence to build-up and allow the unmasking observed in Experiment 1, with no additional enhancement or interference due to jitter/predictability in the precursor. Previous studies have suggested that unmasking due to precursors is initially a “temporal decline in masking” over the first ∼400ms (also observed for sound with no temporal coherence) and then a large threshold decrease (in a subset of participants) after 400ms^39,40^. If there is a build-up benefit to temporal coherence then there was not sufficient time in this experiment for it to manifest (in just the masker) and, hence, it is unlikely we observed an additional enhancement from build-up in the masker, beyond a temporal decline in masking. We observed that enhancement due to temporal coherence could be blocked by manipulating the precursor, as described previously^37-39^. However, this is not “true” interference (i.e. a decrease in threshold relative to a neutral condition) but rather a loss of enhancement. More work would be needed to test whether or to what extent interference of temporal coherence is a simultaneous or a sequential phenomenon.

Other work has also demonstrated the type of interference effects observed here. Grose et al.^38^ presented 4 continuous narrow bands of noise (20 Hz wide centred at 804, 1200, 1747 and 2503Hz) with a temporal structure that was switched from being comodulated (i.e. temporally coherent) to randomly modulated, while trained participants were asked to detect pure tones. The authors found that incoherently modulated precursors/postcursors (both were modified in each trial) produced a small interference with comodulation masking release, elevating thresholds by ∼2dB. Subsequent work by, largely, the same authors^37^ demonstrated a similar effect in highly trained participants (n=4), though with a much larger effect size (∼10dB). Similarly, a large interference (9.55dB) was observed in our study when replacing a comodulated build-up with a temporally jittered one (n=19, minimal training provided). Our work, which only looked at the influence of precursor sounds, also demonstrated an interference attributable to disruption of temporal predictability in the build-up (Fig. 1D), suggesting the interference observed by Grohse and colleagues is largely attributable to build-up rather than postcursor interference.

As mentioned already, in Experiment 1 we found no evidence of the build-up period promoting or interfering with the ability to benefit from for temporal coherence in the masker. In addition, in Experiment 2 removing temporal coherence while keeping temporal predictability in the signal band (TP^+SIG^TC^-^) did not produce significant interference, whereas, removing temporal predictability from the signal band produced the greatest interference. We did observe that enhancement due to temporal coherence could be blocked by manipulating the precursor, as described previously^37-39^. However, this is not “true” interference (i.e. a decrease in threshold relative to a neutral condition) but rather a loss of enhancement. This suggests that either interference/enhancement due to build-up has a different time integration window or that temporal predictability is a relatively instantaneous cue. Frequently in the study of temporal coherence, particularly in streaming, the temporally coherent stimuli is also temporally predictable (during build-up and thereafter), meaning it possesses both temporal predictability and temporal coherence. Our data highlight the need to separate these two cues when studying temporal coherence.

In conclusion, this work supports the idea that build-up is a critical process for shaping subsequent perception. Manipulating temporal coherence and predictability during build-up produces independent changes in enhancement and interference of subsequent tone detection (Experiment 1). In addition, our results suggest that temporal predictability influences perception over longer timescale than temporal coherence alone and can produce large interference in subsequent perception. Further work is needed to understand to what extent stimulus history can influence the use of temporal coherence cues.

## Acknowledgments

This work was funded by a grant from the BBSRC (BB/N001818/1) and a Wellcome Trust/Royal Society Sir Henry Dale Award (098418/Z/12/Z). This research was funded in whole, or in part, by the Wellcome Trust [Grant number 098418/Z/12/Z]. J.S. was also funded by the University of Nottingham, Nottingham Research Fellowship. For the purpose of open access, the author has applied a CC BY public copyright licence to any Author Accepted Manuscript version arising from this submission.

## Author contributions

J.S. designed and coded the experiments. K.C.P. and D.B.M. recruited participants and collected data. J.S. analysed the data. J.S. and J.K.B wrote the manuscript. J.S., K.C.P. and J.K.B. reviewed the manuscript.

## Data availability statement

The dataset from the current study is publicly available through Figshare: https://figshare.com/s/619b2bb486a4d226c3b3

## Additional Information

The authors declare no competing interests.

